# Investigating the causes of stimulus-evoked changes in cone reflectance using a combined adaptive optics SLO-OCT system

**DOI:** 10.1101/2021.10.30.466627

**Authors:** Mehdi Azimipour, Denise Valente, John S. Werner, Robert J. Zawadzki, Ravi S. Jonnal

## Abstract

*In vivo* functional imaging of human photoreceptors is an emerging field, with compelling potential applications in basic science, translational research, and clinical management of ophthalmic disease. Measurements of light-evoked changes in the photoreceptors has been successfully demonstrated using adaptive optics (AO) coherent flood illumination (CFI), AO scanning light ophthalmoscopy (SLO), AO optical coherence tomography (OCT), and full-field OCT with digital AO (dAO). While the optical principles and data processing of these systems differ greatly, and while these differences manifest in the resulting measurements, we believe that the underlying physiological processes involved in each of those techniques are likely the same. AO-CFI and AO-SLO systems are more widely used than OCT systems. However, those systems produce only two-dimensional images and so, less can be said about the anatomical and physiological origins of the observed signal. OCT signal, on the other hand, provides 3D imaging but at a cost of high volume of data, making it impractical to clinical purposes. In light of this, we employed a combined AO-OCT-SLO system–with point-for-point correspondence between the OCT and SLO images–to measure functional responses simultaneously with both and investigate SLO retinal functional biomarkers based on OCT response. The resulting SLO images reveal reflectance changes in the cones which are consistent with those previously reported using AO-CFI and AO-SLO. The resulting OCT volumes show phase changes in the cone outer segment (OS) consistent with those previously reported by us and others. We recapitulate a model of the cone OS previously proposed to explain AO-CFI reflectance changes, and show how this model can be used to predict the signal in AO-SLO. The limitations of the model is also discussed in this manuscript.

## 1 Introduction

Functional assessment of the retina is a critical portion of both clinical ophthalmology and vision research. Several techniques have been developed since the 19th century for measuring the optoretinographic (ORG) responses, such as visual acuity/contrast sensitivity tests, psychophysics and ERG (electroretinogram and multifocal electroretinogram). However, these approaches have important shortcomings such as poor spatial resolution, long duration of tests and, in some cases, being slightly invasive, limiting their application as well as their potential for early stage detection of photoreceptors impairment. The imaging resolution would also play a role to access the degree of success of emerging therapies such as human gene therapy, where new methods are being studied to rescue existing dysfunctional photoreceptors, or in photoreceptors transplantation to restore vision loss in degenerative diseases.

In virtue of that, there is great excitement about new fast and noninvasive approaches with cellular resolution to measure retinal function such as flood-illumination, SLO [2, 3] and OCT [4, 5, 6, 7, 8, 9, 10, 11, 12, 13]. In flood-illumination and SLO the stimulus-evoked response of photoreceptors is detected as changes in reflectance of the cone mosaic using near-infrared imaging light. Some hypothesis to explain this intrinsic signal includes stimulus-evoked outer segment elongation combined with coherent interference within the cone, refractive index change between photoreceptor inner segment and outer segment and also change in scattering properties of outer segment followed by photoisomerization. Meanwhile, the OCT approaches have shown that the photoreceptors outer segments appear to elongate in response to a stimulus.

Overall, the slow dynamic of the functional signal suggests a transduction stage downstream from the initial photoisomerization [3]. In this work, an integrated AO SLO-OCT was employed to tackle some of ambiguities in changes in reflectance of near-infrared imaging followed by delivering visible stimuli.

## 2 Methods

### 2.1 Experimental setup

A schematic of the combined AO-SLO-OCT system[1, 12] is shown in Fig. 1(A). The swept-source OCT employed a Fourier-domain mode-locked (FDML) laser (1063 *±* 39 nm) with an A-scan rate of 1.64 MHz[14], while the SLO light source was a 840*±*25 nm superluminescent diode (S840, Superlum Ltd, Moscow, Russia). For functional experiments, an optical filter centered at 830 nm with 10 nm bandwidth (full-width at half-maximum) was placed in front of the SLO launch, providing a coherence length in the range of 21 µm, assuming a refractive index of 1.4 for the OS. OCT and SLO beams are combined using a dichroic beam splitter (DBS). Apart from the launches of the OCT and SLO beams and their respective detection arms, the beam paths for the two subsystems were identical making the volume and frame acquisition rates the same for the two subsystems and resulting images co-extensive notwithstanding possible small offsets due to transverse chromatic aberration[1].

**Figure 1:**
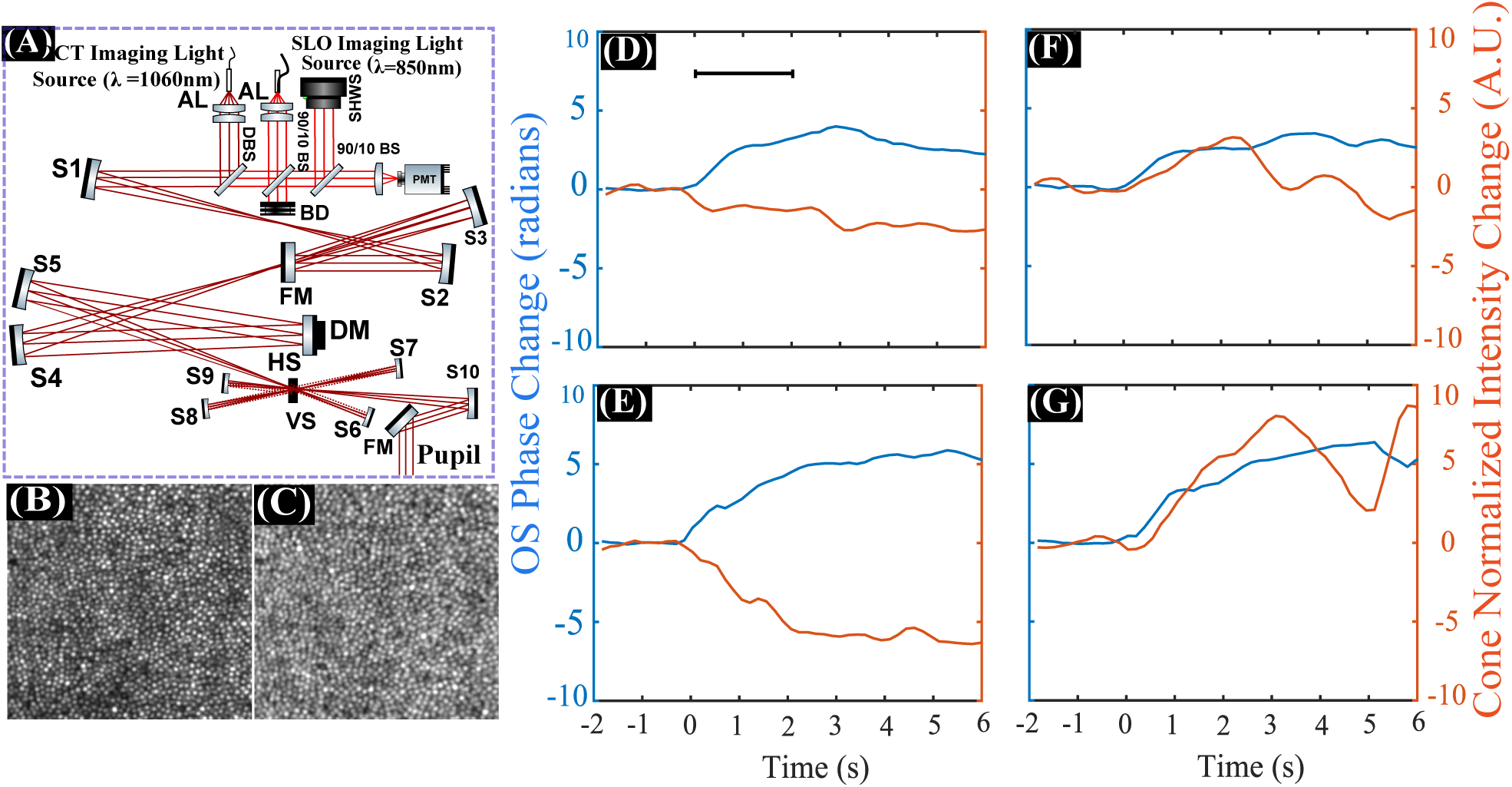
(A) Schematic of the combined AO-SLO-OCT system adapted from [1]. Panels (B) and (C) show examples of an SLO frame and an *en face* projection of the photoreceptors from an OCT volume, respectively, acquired simultaneously at 2° temporal retina (TR). Panels (D)-(G) show cone OS elongation measured using the OCT system (clue curves) and cone standardized reflectance measured using the SLO system (orange curves), in four individual cones in response to a 2 s flash at 555nm with the power of 0.5 µW.The OCT response in all cones shows an increasing after t=0s. Meanwhile, the SLO response presents a wide variety of responses with some going up, down or even oscillating.

**Figure 2:**
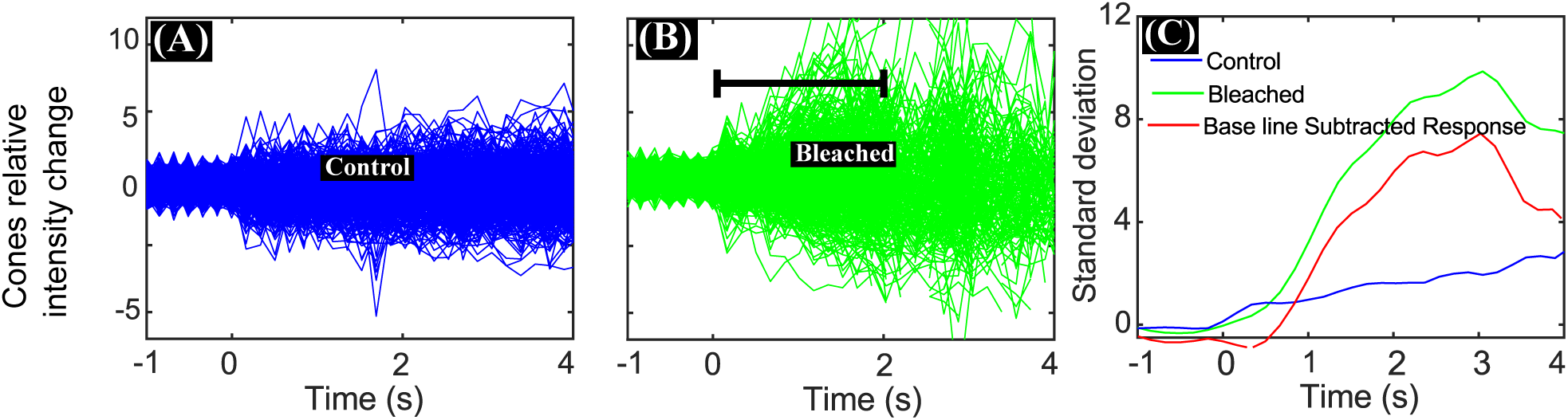
Panels (A) through (C) depict the processing pipeline to extract the aggregate SLO functional signal. First, cones were identified automatically and their standardized reflectance were calculated for the (A) control and (B) bleached (2 s flash at 555nm with the power of 1 µW) images. Then, the spatial standard deviation of standardized reflectance was calculated for each frame, for trials without and with stimuli, as shown in panel (C). Finally, the SLO functional signal was calculated by subtracting the control standard deviation from the stimulus standard deviation.

The system also incorporates a hardware adaptive optics (AO) with a high-speed deformable mirror (DM-97-15; ALPAO SAS, Montbonnot-Saint-Martin, France) and custom-built Shack-Hartmann wavefront sensor (SHWS) consisting of a scientific complementary metal–oxide–semiconductor (sCMOS) camera (Ace acA2040-180 km; Basler AG) and lenslet array (pitch, 500 *µ*m; f 30 mm, Northrop-Grumman Corp, Arlington, Virginia). The SLO light source is used as a beacon to the AO. DM and SHWS operates in close loop using open source software developed in Python/Cython by our lab for real time aberrations correction providing diffraction limited resolution to the system.

A previous implementation of the AO-OCT system alone permitted volumetric images to be acquired at 32 Hz[10], but from the resulting measurements a slower rate of 6 Hz was deemed sufficient for the expected responses to the planned low-power stimulus flashes. This lower rate was achieved by replacing the 5 kHz resonant scanner with a 2 kHz scanner, which resulted in a larger field of view (1.2°) and denser sampling of the retina. The backscattered SLO light from the retina was spatially filtered using a confocal pinhole with a diameter of 1 Airy disk and detected with a photomultiplier tube (PMT). The detection channel of the OCT was identical to what was previously reported elsewhere[10]. The stimulus channel consisted of a green LED combined with a bandpass filter (*λ* = 555nm; Δ*λ* = 20nm), configured in Maxwellian view and coupled into the system with a second dichroic filter in front of the eye. This band was chosen taking into account that it stimulates equally the L and M cones, which make up the overwhelming majority of cones. The stimulus light was modulated with a mechanical shutter in sync with data acquisition in both OCT and SLO subsystems.

### 2.2 Imaging protocol

After obtaining informed consent, three subjects, free of known retinal disease, were imaged. Eyes were dilated and cyclopleged using topical drops of phenylephrine (2.5 %) and tropicamide (1.0 %). A bite bar and a forehead rest were employed to position and stabilize the subject’s pupil during imaging. Subject fixation was guided with a calibrated target. All procedures were in accordance with the tenets of the Declaration of Helsinki and approved by the UC Davis Institutional Review Board.

The time-averaged optical powers of the scanning OCT and SLO beams were respectively 1.8mW and 150 µW at the cornea, together bellow the maximum permissible exposure specified by the 2014 ANSI standard (up to 8 hours) [15]. Prior to each measurement, initial setup consisted of imaging a calibration grid and the subject dark adapted during 5 minutes. After achieving closed-loop diffraction-limited AO correction, a series of 50-100 synchronized OCT volumes and SLO frames were acquired, with a stimulus flash of varying intensity and duration delivered after 2 seconds. Images were acquired at two locations, 2° and 6° temporal retina (TR), where large numbers of both rods and cones are present.

### 2.3 OCT signal processing

Volumetric OCT images were segmented axially and the inner-outer segment (IS/OS), cone outer segment tip (COST) layers were automatically identified and projected [16]. Using images of the calibration grid, sinusoidal distortions were removed using linear interpolation and nearest neighbor interpolation in the SLO and OCT images, respectively. Nearest neighbor interpolation was used for the OCT volumes to prevent phase errors caused by linear interpolation between wrapped phase measurements. The OCT *en face* projections, along with SLO frames, were registered using a strip-based method, producing a trace of the eye movements. Cone photoreceptors were automatically identified and the movement trace was used to track them through the series of 50-100 OCT volumes and SLO frames.

For OCT data analysis, the sub-volumes from the complex OCT data containing individual photoreceptors were analyzed for stimulus-evoked changes. The OCT signal was arranged into M-scans, whose vertical and horizontal dimensions correspond to depth and time, respectively. The magnitude of the OCT signal yields a time-estimate of the cell’s axial reflectance profile while phase difference between COST and IS/OS was used to detect changes in the length of the cone outer segment, as described in detail previously[10]. The volume acquisition rate was not enough to detect the initial rapid contraction reported by others but shows the subsequent elongation in response to the delivered stimulus.

### 2.4 SLO signal processing

The processing pipeline to extract the SLO functional response was identical to one previously reported by Cooper *et al*. [3] and, briefly, consisted of tracing stimulus-evoked reflectance changes using the following the steps: first, cones were identified and tracked as described above. Next, the cone’s time-varying reflectance was standardized by subtracting the average pre-stimulus cone reflectance from the time-varying reflectance of each cone and then dividing by its pre-stimulus standard deviation, such that a cone with non-varying reflectance would have, in theory, a constant reflectance of zero and possible changes in reflectivity would be normalized by the cone individual noise baseline.

While the OCT signal of individual cones shows a monotonical increase shortly after the stimulus, the SLO response of individual cones could present an increasing, decreasing or even oscillation of the stimulus-evoked reflectance, with rate and direction changes essentially random among cones, in accordance with previously reported AO-CFI and AO-SLO.

Finally, the standard deviation across all cones within each frame was calculated to establish the functional change of the ensemble. Since the standard deviation of normalized reflectance changes over time even in absence of stimulus due to noise, a separate control trial with no flash was processed the same way and subtracted from the bleached.

## 3 Simulations

The origin of the observed changes in reflectivity in response to bleaching is still unknown. The AO-CFI functional responses were conjectured to be due to interference, likely between light reflected by the IS/OS and COST [17], and while the coherence length of the light source was not investigated previously in AO-SLO, the observed effect is nearly identical to the former, which suggests a similar role. Other hypothesis suggests that changes in scattering properties of outer segment followed by photoisomerization could be the source of AO-SLO functional response however, one can argument that those changes due to bleaching should follow the same tendency across all photoreceptors, showing either a collective increase or decrease in reflectivity. This hypothesis can also be refuted by analysing individually the reflectivity of inner-to-outer segment junction (ISOS), the cones outer-segment termination (COST) and the retinal pigment epithelium layer (RPE).

AO-OCT imaging reveals directly the phase difference between IS/OS and COST, and can thus be used to simulate and predict the behavior of cones in AO-SLO imaging. To do so, the ORGs collected with the OCT subsystem were fitted to an overdamped harmonic oscillator equation for*t >* 0 (after the stimulus was delivered):

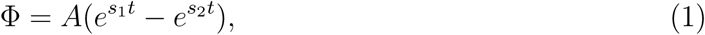

where *A, s*_1_ and *s*_2_ are free parameters. That model is not aimed to describe in details the phototransduction events involved but instead, it qualitatively agrees with the elongation and recovery observed in the ORGs. With that, the expected areal reflectance of the cone considering interference was simulated by the following equation, where an initial phase difference in [0, 2*π*) and initial fractional reflectivities in [0.4, 0.6] were assigned to the IS/OS and COST:

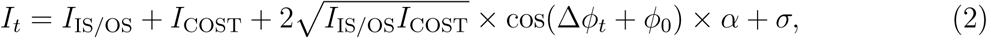

where *I*_ISOS_ and *I*_COST_ are the intensity of IS/OS and COST bands, respectively, *ϕ*_0_ is their initial phase difference and Δ*ϕ*_*t*_ is their phase difference at time point *t*, as measured by the OCT system, *α* is the interference efficiency related to the coherence length of the light source and *σ* is white noise added to the simulation - here, random values from 0 to 0.2. The white noise was necessary to allow a nonzero value for the pre-stimulus standard deviation, used in the SLO pipeline described in session §2.4. A change in the white noise range would impact the amplitude of the SLO functional response but not its shape. The same is also true to the assigned initial fractional reflectivities.

## 4 Results

Figs. 4A and 4B show the AO-SLO image of the cone mosaic, as well as the locations of the automatically identified cones at 6°. As shown in Fig. 4, visible stimuli cause modulation of cone reflectance in the SLO image. After aggregating cone responses as described above [3], we observed that the shape of the resulting aggregate (standard deviation) response was very similar to the elongation observed in the OCT channel. Figs. 4C and 4D show the SLO spatial standard deviation and OCT average OS elongation, respectively, for two different stimulus levels. It is apparent that the shape of these curves is very similar, as shown in Fig. 4E and 4F. We did not have an *a priori* expectation that these would be so similar, and future work will include investigation of this similarity.

**Figure 3:**
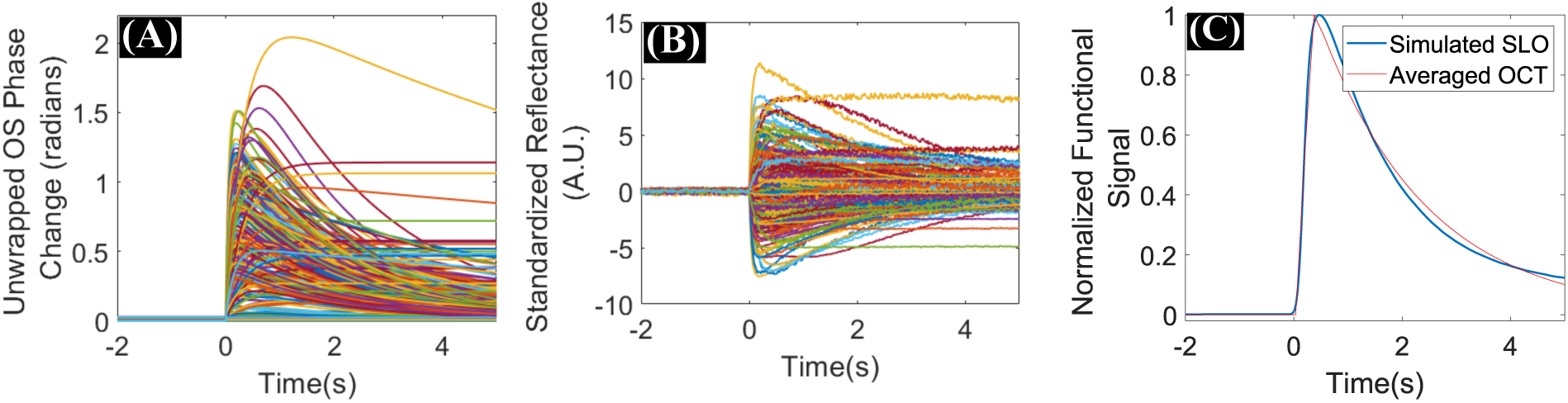
Panels (A) through (C) depict an example of processing pipeline to simulate the aggregated SLO functional signal using AO-OCT. (A) First, RLC function was fitted to the unwrapped phase information. Then, the standardized reflectance was calculated for each cone (B). And finally, the spatial standard deviation of standardized reflectance was calculated as shown in panel (C).

**Figure 4:**
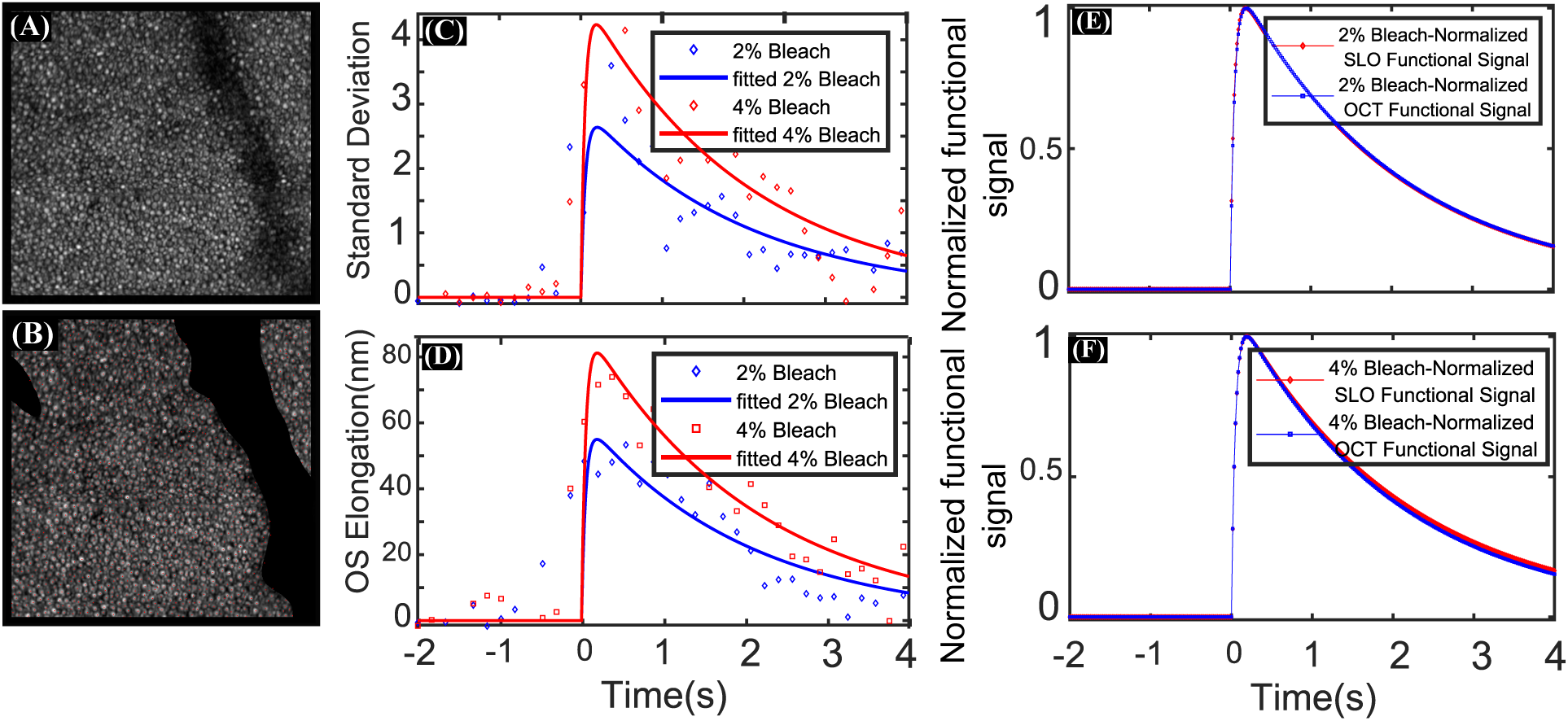
(A) SLO frames acquired at 6° TR, and (B) the coordinates of automatically identified photoreceptors. (C) and (D) show the spatial standard deviation of standardized cone reflectivity–the SLO functional signal–and OS elongation measured with OCT for two flash intensities with duration of 12*ms*. (E) and (F) show normalized SLO and OCT signals plotted together for two flash intensities, showing that the responses to flashes bear significant similarity. This similarity was not expected, owing to the complicated calculation required for the SLO aggregate signal. Nevertheless, it provides confirmation that the SLO signal originates from OS elongation combined with interference within the cone OS.

Figs. 5A and 5B show the OCT elongation and SLO aggregate response for two CW stimuli. In contrast to the flash responses shown in Fig. 5, the OCT and SLO responses here are slightly different for the low bleaching level (Fig. 5C), while the difference is significant for the higher bleaching percentage as shown in Fig. 5D. The OCT functional signal shows a monotonically increasing pattern for the higher bleaching level, while SLO signal shows an oscillating pattern that increases at the beginning and, while the light was still on, decreases after a few seconds and then increases again. To investigate the unexpected behavior of the SLO signal, the model introduced in section 2.4 was employed to predict the SLO signal based on cone OS elongation. First, a few hundred cones were selected from the OCT volumes and their stimulus-evoked elongation was measured. Eq. (2) allowed us to predict changes in their reflectance resulting from elongation, and these changes were used to simulate their behavior upon SLO imaging. The simulated SLO responses and observed responses do bear significant similarity, as shown in Figs. 5E and 5F, which suggests that the source of SLO reflectance changes is coherent interference within the cone OS.

**Figure 5:**
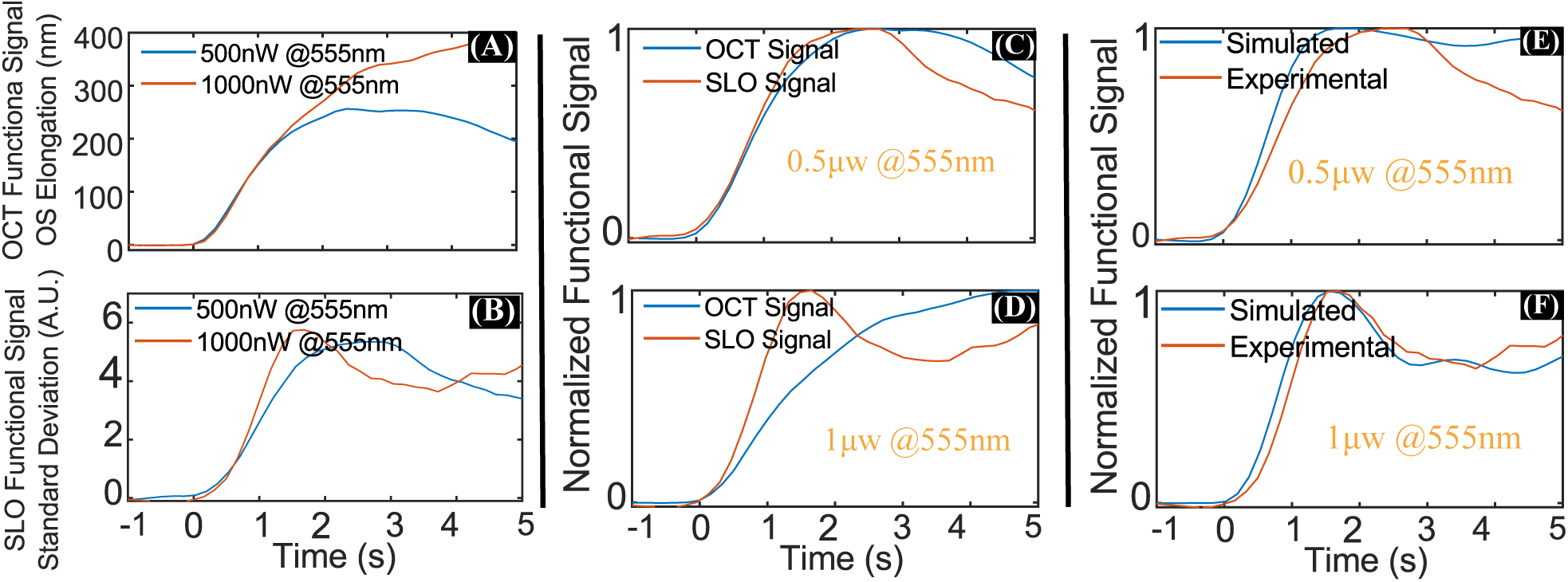
Functional OCT and SLO signals were recorded simultaneously during two different CW stimulus exposures and plotted separately in panels (A) and (B), respectively. The OCT functional signal shows a larger, monotonically increasing pattern for the higher bleaching level, unlike SLO functional signal, which shows an oscillating pattern that increases at the beginning and, while the light was still on, decreases after a few seconds and then increases again. The normalized OCT and SLO signals for these two experiments are shown in panels (C) and (D), respectively. To further investigate the unexpected behavior of the SLO signal, the model introduced in section 2.4 was employed to predict the SLO signal based on cone OS elongation. The results are shown in panels (E) and (F). The simulation results are in a good agreement with the experimental findings and further suggest that the underlying SLO signal originate from coherent interference between retinal bands.

**Figure 6:**
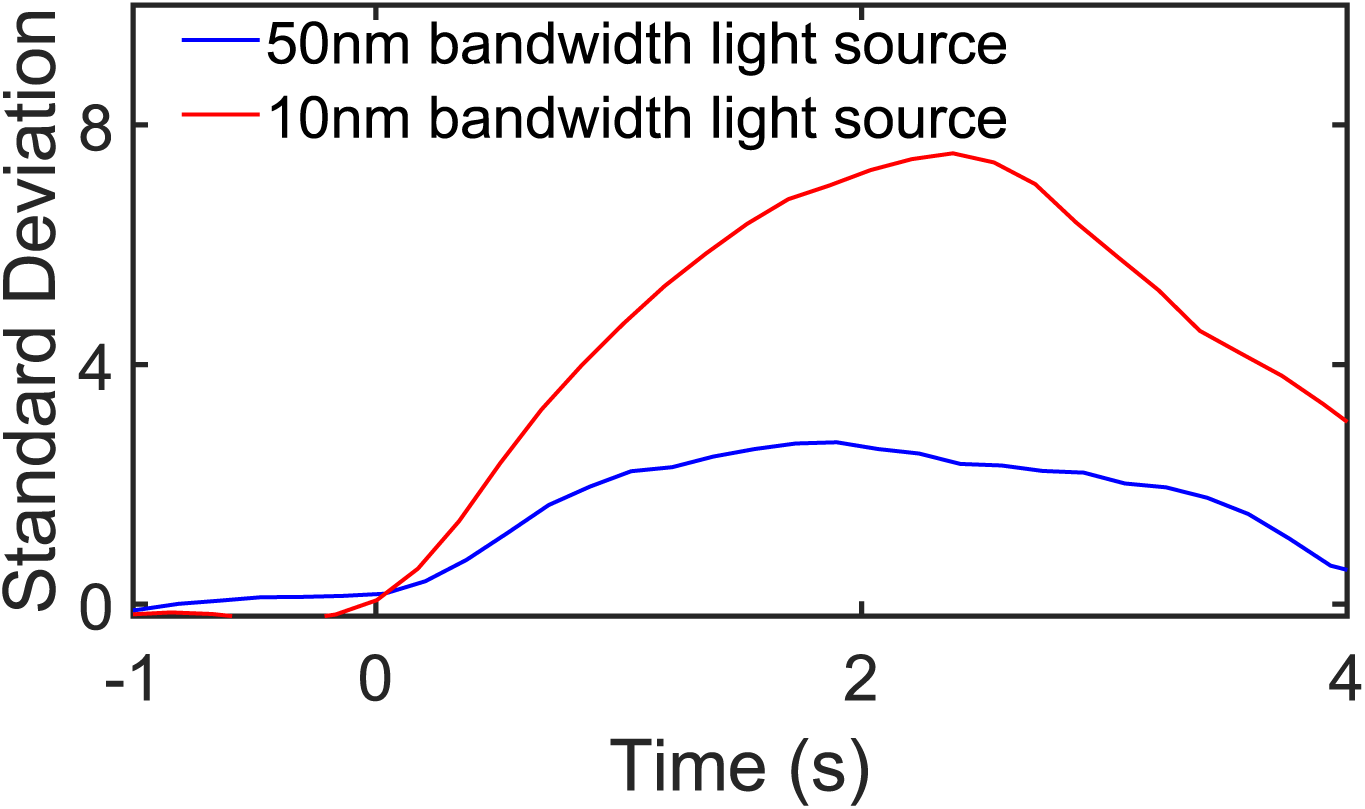
Functional SLO signals were captured using an imaging light source with two different optical bandwidth of 50nm and 20nm. As expected, by increasing the bandwidth of the light source (lowering the coherence length), the amplitude of the SLO functional was decreased which is the consequence of lower interference efficiency between the backreflected light from cones axial bands, especially IS/OS and COST.

To further investigate this hypothesis, we performed identical experiments with the full 50 nm bandwidth of the SLO source and consequently shorter temporal coherence length. The resulting images showed reduced responses, but we are not sure if this residual response remains because the OS length falls sufficiently within the coherence envelope as to generate an appreciable interference component, or if it is due to the presence of additional reflections within the OS[18]. This question will be a topic of future investigation.

## 5 Conclusions

Our results suggest that the functional AO-SLO signal may be estimated using AO-OCT measurements and the former may not provide any additional complementary knowledge. Moreover, given that at present the functional SLO signal must be aggregated over hundreds or thousands of cells, its resolution is similar to that of electrophysiological and psychophysical approaches described above, and much lower than the cellular resolution afforded by AO-OCT. Lastly, the dynamic range of the AO-SLO signal is likely limited by the fact that it is an oscillating signal which only increases inter-cell variance initially. Nevertheless, AO-SLO offers the relative advantages of simplicity, smaller data volumes, and wider deployment. Thus continued investigation of the relationship between these signals using multimodal AO systems may be of great value, and may lead to faster and broader investigation of questions about basic visual processes and disease-related dysfunction.

## Notes

### Competing Interest Statement

The authors have declared no competing interest.

